# RnaBench: A Comprehensive Library for *In Silico* RNA Modelling

**DOI:** 10.1101/2024.01.09.574794

**Authors:** Frederic Runge, Karim Farid, Jörg K.H. Franke, Frank Hutter

## Abstract

RNA is a crucial regulator in living organisms and malfunctions can lead to severe diseases. To explore RNA-based therapeutics and applications, computational structure prediction and design approaches play a vital role. Among these approaches, deep learning (DL) algorithms show great promise. However, the adoption of DL methods in the RNA community is limited due to various challenges. DL practitioners often underestimate data homologies, causing skepticism in the field. Additionally, the absence of standardized benchmarks hampers result comparison, while tackling low-level tasks requires significant effort. Moreover, assessing performance and visualizing results prove to be non-trivial and task-dependent. To address these obstacles, we introduce RnaBench (RnB), an open-source RNA library designed specifically for the development of deep learning algorithms that mitigate the challenges during data generation, evaluation, and visualization. It provides meticulously curated homology-aware RNA datasets and standardized RNA benchmarks, including a pioneering RNA design benchmark suite featuring a novel real-world RNA design problem. Furthermore, RnB offers baseline algorithms, both existing and novel performance measures, as well as data utilities and a comprehensive visualization module, all accessible through a user-friendly interface. By leveraging RnB, DL practitioners can rapidly develop innovative algorithms, potentially revolutionizing the field of computational RNA research.

## 1 Introduction

RNA molecules play critical roles in the regulation of various biological processes, such as cellular differentiation and development [Morris and Mattick, 2014]. These functions are intricately tied to the hierarchical formation of RNA structures [Tinoco Jr and Bustamante, 1999]. Initially, rapid intra-molecular nucleotide interactions lead to the creation of local geometries, known as the *secondary structure*, which governs the interaction regions with other cellular components [Gandhi et al., 2018]. Consequently, this secondary structure guides the subsequent formation of the ultimate 3-dimensional shape, referred to as the *tertiary structure* [Tinoco Jr and Bustamante, 1999]. Therefore, accurately predicting the secondary structure of RNA and designing RNA sequences that fold into desired structures are two fundamental challenges in computational biology [Bonnet et al., 2020], holding significant implications in fields such as medicine, synthetic biology, and biotechnology.

The potential of deep learning (DL) algorithms in biological structure prediction tasks is undeniable. We have witnessed significant impacts of DL methods in the field of protein structure prediction, as evidenced by notable advancements achieved by recent studies [Jumper et al., 2021, Lin et al., 2022]. Additionally, the field of cheminformatics has experienced an explosion of generative DL-based algorithms, due to their potential in the exploration of the chemical space and the prediction of novel compounds [Engkvist et al., 2021]. Given these successes, it is reasonable to expect that accurate DL-based methods could have similar transformative effects on RNA computational biology. However, it is puzzling that they have not received the same level of attention and recognition in this particular domain.

The initial success of the DL algorithm, *SPOT-RNA* [Singh et al., 2019], paved the way for numerous DL-based approaches that have achieved state-of-the-art performance in RNA secondary structure prediction [Zhang et al., 2019, Rezaur Rahman Chowdhury et al., 2019, Chen et al., 2020, Singh et al., 2021, Fu et al., 2022, Saman Booy et al., 2022, Wayment-Steele et al., 2022, Jung et al., Chen et al., 2022, Franke et al., 2022, Chen and Chan, 2023]. However, subsequent investigations revealed that the observed performance gains were potentially driven primarily by learning homologies within the data [Flamm et al., 2021, Szikszai et al., 2022], and overfitting became a concern [Sato et al., 2021]. Consequently, skepticism regarding DL methods has arisen within the research community. To regain trust and establish DL methods in the field, it is crucial to develop appropriate data pipelines and standardized datasets that properly account for biological homologies. However, this requirement often poses challenges for non-domain experts.

RNA design refers to the reverse problem in which the objective is to identify an RNA sequence that can fold into a given target structure [Zuker and Stiegler, 1981, Hofacker et al., 1994]. Various approaches have been proposed to tackle this challenge, including stochastic local search [Andronescu et al., 2004], constraint programming [Garcia-Martin et al., 2013], evolutionary algorithms [Esmaili-Taheri et al., 2014], ant-colony optimization [Kleinkauf et al., 2015], and reinforcement learning [Eastman et al., 2018, Runge et al., 2019]. However, the lack of standardized benchmark datasets poses a significant hurdle for the development and evaluation of DL methods in this field. Currently, the only widely recognized benchmark dataset available is the Eterna100 testset [Anderson-Lee et al., 2016], which consists of 100 synthetic samples lacking any training data. Furthermore, evaluation protocols differ across approaches that report on this test set, further complicating the comparison and assessment of different methods. Consequently, these challenges have impeded the progress and adoption of DL methods in RNA design, leading to their scarcity in the field.

To address the existing challenges and facilitate the entry of DL methods into the field of RNA structure prediction and design, we introduce RnaBench (RnB)1 — a comprehensive RNA benchmark library. RnB has been developed with the aim of providing high-quality datasets, standardized evaluation protocols, and comprehensive analysis, to tackle the important and intriguing challenges of RNA structure prediction and design in computational biology. The key contributions of RnB are as follows:

- RnB includes three benchmarks for RNA secondary structure prediction, accompanied by standardized evaluation protocols (Section 3.1).
- RnB offers meticulously curated RNA datasets that take into account biological homology, ensuring that DL algorithms are trained on relevant and representative data (Section 3.1.1).
- We introduce novel performance measures that address previously untracked aspects of algorithm performance, allowing for more comprehensive evaluations (Section 3.1.2).
- RnB presents the first RNA design benchmark suite, complete with standardized evaluation protocols, providing developers with a means to assess their algorithms in the context of RNA design (Section 3.2).
- We develop a novel RNA design benchmark based on a real-world Riboswitch design problem, enabling the application of generative DL algorithms to the RNA design problem, akin to approaches in cheminformatics (Section 3.3).
- RnB incorporates a comprehensive visualization module that facilitates the analysis and interpretation of predictions (Section 3.4).

## 2 Related Work

Existing RNA libraries [Lorenz et al., 2011, 2016, Reuter and Mathews, 2010] primarily serve the needs of biological researchers and experimental practitioners, providing utilities for tasks such as file conversions, specific computational prediction methods, and analysis. However, these libraries lack curated data, data pipelines, and tools for comparing algorithm performance.

While various databases house large amounts of biological data [Berman et al., 2000, Griffiths-Jones et al., 2003, Danaee et al., 2018, wwp, 2019, Consortium, 2020] and software tools exist that provide interfaces to interact with such databases [Cock et al., 2009], the careful curation of data remains a responsibility placed on the users, demanding substantial domain knowledge. Currently, the only data processing pipeline geared towards supporting algorithm development directly is RNAcmap [Zhang et al., 2021], which facilitates the automatic search for homologous sequences through direct coupling analysis. However, data curation still relies on the users. In contrast, RnaBench is specifically designed to address these limitations by providing comprehensive pipelines and standardized datasets. Our focus is to enable non-domain experts and algorithm developers to readily apply algorithms to RNA structure prediction and design tasks, reducing the effort required for data curation and promoting out-of-the-box applications.

Regarding benchmarking, the only current available benchmark for RNA secondary structure prediction is EternaBench [Wayment-Steele et al., 2022], which relies on synthetic RNA data derived from human design approaches on the Eterna crowdsourcing platform [Lee et al., 2014]. In RnaBench, we shift the focus to experimentally derived data. Additionally, EternaBench assesses the prediction of structure ensembles represented as base pair probabilities. While this is generally desirable due to the dynamic nature of RNA [Ganser et al., 2019], recent research has shown that appropriate DL algorithms can learn structure distributions from single structure data [Franke et al., 2022]. Therefore, in RnaBench, we concentrate on the more commonly approached problem of single secondary structure prediction.

For RNA design, the widely used Eterna100 benchmark dataset [Anderson-Lee et al., 2016] is also based on synthetic data from the Eterna platform. This dataset comprises only 100 samples without any training and validation data. Further, the commonly used version one of this benchmark includes tasks that have been proven to be unsolvable with certain folding algorithms [Koodli et al., 2021], which has been resolved in the less commonly known version two of the Eterna100 testset. RnaBench overcomes this limitation by including multiple test sets, training and validation data, and a range of folding algorithms to address these challenges effectively.

## 3 RnaBench

In this section, we introduce RnaBench (RnB), a comprehensive open-source library dedicated to *in silico* RNA data modeling. RnB offers a diverse range of benchmarks that encompass RNA secondary structure prediction (Section 3.1), RNA design (Section 3.2), and a novel Riboswitch design benchmark that addresses a real-world RNA design challenge, thereby pioneering generative RNA design with specific properties (Section 3.3). Complementing the benchmarks, RnB provides robust data utilities and advanced visualization tools (Section 3.4), empowering algorithm developers to thoroughly evaluate and analyze the performance of their methods. We note that we use several publicly available datasets for our benchmarks in RnaBench. Table 1 provides an overview of the data. Consistent with the literature [Singh et al., 2019], we use a length cutoff at 500 nucleotides for all datasets. More details about each dataset can be found in Appendix B.

**Table 1:** Overview of the benchmarks of RnaBench.

| Benchmark | Task | Set | # Samples | Min – Max Length |
| --- | --- | --- | --- | --- |
| Intra-family | Structure Prediction | test | 1458 | 22 – 499 |
|  |  | valid | 1247 | 33 – 497 |
|  |  | train | 58271 | 23 – 500 |
| Inter-family | Structure Prediction | test | 153 | 33 – 189 |
|  |  | valid | 17 | 33 – 159 |
|  |  | train | 56621 | 12 – 500 |
|  |  | fine-tune | 2766 | 12 – 500 |
| Biophysical Model | Structure Prediction | test | 3344 | 37 – 182 |
|  |  | valid | 2720 | 34 – 160 |
|  |  | train | 615204 | 24 – 500 |
| Inverse RNA Folding | Design | test | 1379 | 12 – 497 |
|  |  | valid | 402 | 16 – 500 |
|  |  | train | 66772 | 12 – 500 |
| Constrained Design | Design | test | 315 | 21 – 274 |
|  |  | valid | 89 | 16 – 447 |
|  |  | train | 77148 | 12 – 500 |
| Riboswitch Design | Generative Design | train | 685109 | 67 – 91 |

### 3.1 RNA Secondary Structure Prediction Benchmark

RNA secondary structure prediction algorithms can be roughly divided into two classes: (1) *de novo* prediction methods that seek to predict a structure directly from the sequence of nucleotides and (2) *homology modeling* methods that require a set of homologous RNA sequences for their predictions [Singh et al., 2021], called an RNA family. Predictions can then be applied either within given families (intra-family predictions) or across different families (inter-family prediction). *De novo* prediction methods are typically preferred, particularly when working with novel RNAs, as searching for homologous sequences can be time-consuming and challenging, and in many cases, no family information is available [Szikszai et al., 2022]. By emphasizing *de novo* prediction methods in our benchmark for secondary structure prediction, we aim to provide a comprehensive evaluation of algorithms that can tackle the challenges of RNA structure prediction without relying on prior knowledge of homologous sequences or families. Our secondary structure prediction benchmark provides three tasks; Intra-family prediction, inter-family prediction, as well as learning of a simplified biophysical model as proposed by Flamm et al. [2021]. The latter allows us to assess whether a folding algorithm is generally capable of learning the underlying biophysical dynamics of RNA secondary structure formation.

#### 3.1.1 Data

##### Test Data

For assessing intra- and inter-family prediction performance, we use the commonly used validation and test sets VL0, VL1, TS0, TS1, TS2, TS3, and TS-hard provided by Singh et al. [2019] and Singh et al. [2021]. Except for TS0 and VL0 which we only use for intra-family predictions, these datasets are derived from experimentally supported 3D RNA structures from the RCSB Protein Data Bank (PDB) [wwp, 2019]. For the benchmark task of learning a biophysical model, we further use RNAFold [Lorenz et al., 2011] and the RNA Family Database (Rfam) [Griffiths-Jones et al., 2003] to generate a large set of training, validation, and test data in a family-based fashion to account for homologies as described below.

##### Initial Training Data Pool

Before processing the data for the respective tasks of intra- and inter-family predictions, we collect a large initial pool of publicly available RNA sequence and structure pairs from the following sources: the bpRNA-1m meta-database [Danaee et al., 2018], the ArchiveII [Sloma and Mathews, 2016] and RNAStrAlign [Tan et al., 2017] datasets provided by Chen et al. [2020], all data from RNA-Strand [Andronescu et al., 2008], all datasets used in Sato et al. [2021], the Rfam-learn dataset provided by Runge et al. [2019], all datasets used in [Kleinkauf et al., 2015], the dataset provided by Taneda [2010], the Eterna100 datasets version 1 and 2 [Anderson-Lee et al., 2016, Koodli et al., 2021] as well as all RNA containing data from PDB [wwp, 2019], downloaded in May 2022. Secondary structures for PDB samples were derived from the 3D structure information using DSSR [Lu et al., 2015], which was also used in previous work [Singh et al., 2019]. After removing duplicates and dropping samples with less than 90% canonical nucleotides (A, C, G, U), our initial training pool encompasses 111295 samples. The final pool is provided with RnaBench and subsequently used to derive training data for our benchmarks.

##### Intra Family Data

For building data for intra-family predictions, we follow the pipeline of Singh et al. [2019]. In particular, we resolve IUPAC nucleotides [Johnson, 2010] by mapping each nucleotide to either A, C, G, U, or N, where N represents the standard wildcard used in the field to denote unknown nucleotides. We then remove sequence similarity between the test set, the validation set and the training pool at an 80% similarity threshold using CD-HIT [Fu et al., 2012]. To further remove similarities, we drop samples from the training and validation set that show any homology with the test samples, using BLAST search [Altschul et al., 1997] with a large e-value of 10.

##### Inter Family Data

Our inter-family data pipeline is set on top of the intra-family data pipeline to further account for homologies in the data. We develop a novel pipeline inspired by the dataset curation of Singh et al. [2021] for TS-hard and the data pipelines of Rfam. In particular, we first apply the intra-family pipeline to remove sequence similarities between training, validation (we only use VL1 here), and test data. For each sample in the test sets, TS1, TS2, TS3, and TS-hard, we use BLAST search to find homologous sequences, using NCBI’s nt database as a reference. We then use LocARNA-P [Will et al., 2012] to derive multiple sequence alignments based on the sequence and structure similarity of the homologs for each sample in the test set. We build covariance models from the alignments using Infernal [Nawrocki and Eddy, 2013]. The covariance models are used to remove sequences from the train and validation sets that belong to one of the covariance models at an e-value of 0.1.

##### Biophysical Model Data

To generate the datasets for learning a simplified biophysical model, we sample a large number of sequences from all families of the Rfam database with a covariance model of a maximum CLEN of 500 nucleotides. We use all sequences from covariance models with CLEN between 200 and 500 nucleotides and add twice the amount of shorter sequences from covariance models with a CLEN of less than 200 nucleotides to account for the length distribution of the other test sets. All sequences are folded with RNAFold [Lorenz et al., 2011]. From all families in the dataset, we sample 30 and 25 test and validation families at random and kept all samples of other families in the initial pool for training. For compatibility with all other benchmark sets, we additionally remove the similarity between the biophysical model data and all other test and validation sets as described for the intra- and inter-family data pipelines.

##### Inter Family Fine Tuning Data

We provide a training dataset of experimental data only that is consistent with the intra- and inter-family benchmark sets as well.

#### 3.1.2 Evaluation

For evaluation, we employ commonly used and novel performance measures. The commonly used performance measures for RNA secondary structure prediction are *Precision, Recall, F1 Score, Shifted F1 Score*, and *Matthews Correlation Coefficient* (MCC). Further, we use the *Weisfeiler-Lehman (WL) Graph Kernel* [Shervashidze et al., 2011] *for comparing the structure graphs. To our knowledge, the WL kernel was never used to assess the performance of RNA secondary structure prediction algorithms before but shows some advantages over commonly used measures such as F1 Score and MCC [****?****]. We also report the percentage of completely Solved Structures* since this is the actual goal of single secondary structure prediction. RnaBench further provides per-sample timing, while the mean runtime is provided as an additional performance measure. We provide a more detailed description of every performance measure in Appendix C.

#### 3.1.3 Baselines

We provide the following commonly used baseline algorithms: RNAFold [Lorenz et al., 2011], LinearFold-C and LinearFold-V [Huang et al., 2019], ContraFold [Do et al., 2006], IpKnot [Sato et al., 2011], pKiss [Janssen and Giegerich, 2015], and the DL approaches SPOT-RNA [Singh et al., 2019], MXFold2 [Sato et al., 2021], UFold [Fu et al., 2022], the ProbTransformer [Franke et al., 2022], and the RNAformer [Franke et al., 2023]. We detail all baselines in Appendix D.

### 3.2 RNA Design

For the RNA design benchmark, we propose two different tasks, inverse RNA folding, and constrained inverse RNA folding. While the former considers the problem of finding an RNA sequence that folds into a given secondary structure, the latter considers additional positional constraints in the designed sequence. For both tasks, we provide datasets with and without pseudoknotted [Staple and Butcher, 2005] (non-nested) RNAs and support RNA design for a given G and C nucleotide ratio (GC-content), since the GC-content could have a strong impact on the function of an RNA [Isaacs et al., 2006, Chan et al., 2009, Wang et al., 2014].

#### 3.2.1 Data

For the inverse RNA folding benchmark, we use the Eterna100 test sets version 1 and 2 [Anderson-Lee et al., 2016, Koodli et al., 2021], the Rfam-learn test set [Runge et al., 2019], the test set provided by Taneda [2010], and the test set derived from RNA-Strand provided by Kleinkauf et al. [2015], as well as all pseudoknot containing samples of the ArchiveII [Sloma and Mathews, 2016] dataset as provided by Chen et al. [2020]. For constrained inverse RNA folding, we use the test sets derived from the Rfam and the PseudoBase++ [Taufer et al., 2009] databases as provided by Kleinkauf et al. [2015]. For all datasets, we ensure that there is no overlap between the structures of the test sets and our initial training pool and sampled validation sets of appropriate sizes for each benchmark from the remaining training data uniformly at random.

#### 3.2.2 Evaluation

We evaluate RNA design using the performance measures described in Section 3.1.2, in addition to the traditional number of solved tasks within a time limit. This approach enables us to assess algorithms for optimality, standardize the evaluation process, and facilitate comprehensive analysis.

#### 3.2.3 Baselines

We provide RNAInverse [Hofacker et al., 1994, Lorenz et al., 2011] and a deterministic algorithm as baselines. The deterministic algorithm places an A at sites of unpaired nucleotides and the most stable base pair G-C at positions that should pair in the structure. While remarkably simple, this algorithm is already capable of solving some of the structures (Section 4) and can be considered as implementing the trivial solution.

### 3.3 Riboswitch Design Benchmark

Riboswitches [Mironov et al., 2002] are regulatory RNA elements, typically located in the 5^*′*^ untranslated region of messenger RNAs (mRNAs), that can specifically bind certain metabolites (ligand) to alter gene expression. They consist of an *aptamer* for recognizing the ligand, closely connected to an *expression platform* that couples ligand binding with gene regulation. The binding signal is transduced by a conformational change of the switch. Riboswitches are typically found in bacteria but can also be artificially constructed. Since aptamers are capable of binding nearly every molecular and supramolecular target [Vorobyeva et al., 2018], Riboswitches have become interesting tools for different applications including modulation of cellular functions [Hallberg et al., 2017] or the development of biosensors [Findeiß et al., 2017]. For our novel Riboswitch design benchmark, we reimplement an existing protocol for the computational design of transcriptional activating theophylline dependent riboswitches proposed by Wachsmuth et al. [2012]. Generally, we seek to enable evaluations of generative RNA design algorithms similar to approaches in the field of generative design of small organic molecules [Franke et al., 2022]. In particular, the task of an algorithm is to learn the distribution of Riboswitch sequences (and structures) from a large training set and to generate a set of similar but different sequences. These sequences are then folded and evaluated. We further provide specific properties to allow for conditional generative design. To the best of our knowledge, this is the first approach to establish a benchmark for generative algorithms in the field of RNA computational biology.

#### 3.3.1 Data

We generated 685,109 unique sequences and 39,671 distinct structures using the original design pipeline by Wachsmuth et al. (2012), meeting the benchmark’s design criteria. Our dataset includes GC-content for all sequences and energies, enabling property-based conditional generative design.

#### 3.3.2 Evaluation

For evaluation, we reimplement the original protocol of Wachsmuth et al. [2012]. Each of the designed candidate sequences is folded using RNAFold and verified for the existence of different sequence and structure features proposed by Wachsmuth et al. [2012]. A candidate that passes all criteria counts as valid and we report the number of unique valid candidates and the fraction of unique structures in the final pool of valid candidates. More details about the evaluation can be found in Appendix C. Additionally, we allow analyzing predictions with respect to different measures that assess the learning behavior from a distribution learning point of view.

##### Novelty

Effective generative models generate novel RNA sequences that are not present in the training set, ensuring broad coverage, mitigating overfitting, and avoiding memorization of the training data. Therefore, novelty is a key criterion in assessing RNA algorithms, as the limited training data represents only a fraction of the expansive RNA space. In our evaluation, we assess the novelty of two key features: sequence and structure. For sequence novelty, we directly compare the generated RNA sequences to those in the training set. This demonstrates the models’ ability to explore the vast landscape of RNA sequences and produce diverse outputs. To evaluate structure novelty, we focus on essential RNA structural elements like stem and hairpin lengths, as well as the size of inner loops. We also examine the novelty of structure pairs to understand the unique relationships and interactions captured by the generative models. This analysis helps us evaluate the model’s capacity to explore and generate previously unseen structural motifs, contributing to the discovery of novel RNA patterns. We use two measures to assess the novelty, IOU (Intersection over Union) complement and Novelty.

##### Diversity

We assess models based on their capability to generate a diverse set of distinct RNA structures, penalizing repetitions and emphasizing the generation of unique and varied RNAs.

Diversity differs from novelty in that it quantifies the variation between elements within the generated dataset. To measure this variation, we employ different metrics such as Hamming distance for sequences. The Hamming distance measures the dissimilarity between RNA sequences while disregarding the distances between sequences of different lengths. For assessing diversity in structure pairs, we employ the Weisfeiler-Lehman (WL) Graph Kernel. Diversity and Diameter are two key measures used to assess the diversity of generated RNA structures. By incorporating these measures, our evaluation provides a comprehensive analysis of model diversity, ensuring the generation of a wide range of unique RNA structures. This approach enables us to explore the intricate landscape of RNA secondary structures and advance the field of RNA design.

##### KL Divergence

In RNA, we utilize the Kullback-Leibler (KL) divergence to evaluate the realism and diversity of the generated RNA structures. It compares the distributions of key features, such as structural motifs (e.g., hairpin lengths and counts) and sequence features (e.g., GC-content), between the original and generated datasets. This analysis provides valuable insights into the extent to which the generated sequences deviate from the characteristics of the original data, refer to table 3.

Table 2 shows an example of the distribution learning metrics for a subset of the dataset of the riboswitch design benchmark. We note, however, that the evaluation regarding distribution learning metrics is not restricted to the riboswitch design benchmark, but could be used during evaluations of the general RNA design benchmarks and the RNA secondary structure prediction benchmarks as well. In the latter case, these metrics could e.g. inform about common failure cases of a model.

**Table 2:** Riboswitch generation evaluation on a 50k random subset of samples of the original dataset.

| Feature | Novelty |  | Diversity <sup>†</sup> |  |
| --- | --- | --- | --- | --- |
|  | IoU Complement | Novelty | Diversity | Diameter |
| Stem length | 1.0 | 1.0 | 1.5 | 5.75 |
| Hairpin length | 0.988 | 0.4 | 1.99 | 8.5 |
| Inner loop 1<sup>st</sup> length | 0.988 | 0.402 | 1.98 | 9.0 |
| Inner loop 2<sup>nd</sup> length | 0.995 | 0.725 | 0.953 | 8.0 |
| Sequence | 1.0 | 1.0 | 0.053 | 0.075 |
| Structure pairs | 1.0 | 1.0 | 0.432 | 1.0 |

<sup>†</sup>: To assess diversity, we employ a specific distance measure for each feature. We use the Hamming distance to evaluate the sequence, utilize WL for evaluating the structure pairs, and employ the Euclidean distance measure for the remaining features.

#### 3.3.3 Baselines

We use the proposed design procedure of Wachsmuth et al. [2012] as a baseline. Originally, Wachsmuth et al. [2012] constructed riboswitch candidates from (1) the TCT8-4 theophylline aptamer sequence and structure, (2) a spacer sequence of 6 to 20 nucleotides (nt), (3) a sequence of 10nt to 21nt complementary to the 3^*′*^-end of the aptamer, and (4) a U-stretch of 8nt at the 3^*′*^-end of the construct. To generate candidates, Wachsmuth et al. [2012] designed a large library of random sequences for the spacer region (6-20nt) and a library of sequences complementary to the 3^*′*^-end of the aptamer (10-21nt). From these sets, randomly sampled sequences were combined with the aptamer and the 8-U-stretch.

### 3.4 Visualization and Utilities

We provide a set of additional utilities with RnaBench. In particular, we implement different low-level tasks, like converting between different RNA secondary structure representations and reading different file formats, as well as data loading utilities for all datasets to facilitate the application of models with different requirements. We further provide a comprehensive visualization module that supports the plotting of RNA structures, as well as a detailed analysis of the predictions. All plots in the paper are directly generated from within RnaBench. As an additional feature, we support RNA 3D data by providing a pipeline for downloading, parsing, and loading of cleaned RNA 3D Structures. In particular, we use the RNAsolo [Adamczyk et al., 2022] repository to obtain non-redundant 3D RNA structures at different resolution thresholds and provide data utilities for parsing and loading the data. Finally, we provide a simple RMSD calculation to assess predictions of 3D structures. We believe that this feature of RnaBench might be particularly useful to train algorithms for the Critical Assessment of Techniques for Protein Structure Prediction (CASP) competition, which included a specific track on RNA structure prediction this year for the first time.

### 3.5 API

RNA secondary structures can be represented in different ways, either as strings represented in dot-bracket format [Hofacker et al., 1994], as lists of pairs, or as binary adjacency matrices or contact maps with dimensions *L × L*, where *L* is the length of the sequence. Since the dot-bracket notation cannot represent all nucleotide interactions and a matrix representation does not contain information about the nesting of a structure, we use a list of pairs representation for RnaBench. More precisely, we represent each nucleotide interaction in the structure as a triple of the two pairing positions and a page-number [Danaee et al., 2018] that describes the level of nesting of the pair. To interact with RnaBench, the user simply defines a function that wraps the model predictions. We detail the different RNA representations as well as the API in Appendices B and E, respectively.

## 4 Experiments

In this section, we demonstrate the benefits of RnaBench with three experiments. We assess the performance of the two DL baselines compared to traditional methods on the inter-family prediction benchmark and evaluate our deterministic baseline for RNA design on the inverse RNA folding benchmark across all available folding algorithms. Finally, we plot an example of a 5SrRNA of *Drosophila melanogaster* (RNAcentral Id: URS00003B4856_7227) to show the flaws of commonly used performance measures, F1 Score, MCC, and shifted F1 Score, as well as the benefits of our new performance measure, the Weisfeiler-Lehman graph kernel. We show additional results, e.g. an evaluation of the Riboswitch design baseline in Appendix C.

### RNA Secondary Structure Prediction

The results for the evaluation on the inter-family benchmark are shown in Figure 1. The DL baselines generally outperform all other baselines, with SPOT-RNA showing the best overall performance. However, the traditional baselines can only predict the most common, so-called canonical base pairs (A-U, G-C, G-U, and vice-versa), and show superior performance on tasks that only contain these types of pairs (Figure 1, right). While the two DL methods thus seem to improve on non-canonical base pair predictions, they fail to achieve the same performance on the most common base pairs, compared to more traditional methods.

**Figure 1.**
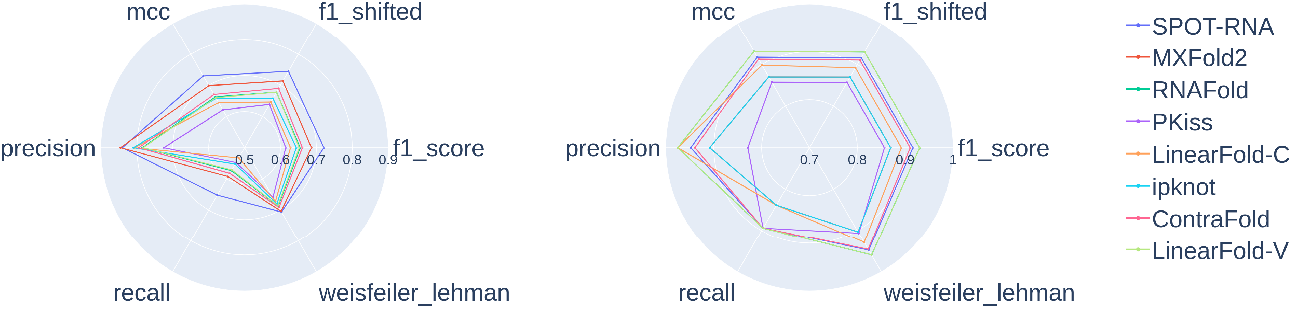
Performance of secondary structure prediction baselines on the inter-family prediction benchmark. (Left) Performance across different performance measures on all types of base pairs. (Right) Performance on canonical base pairs only.

### RNA Design

It was recently shown that the performance of neural network-based approaches for RNA design decreases when changing the folding algorithm [Koodli et al., 2021]. The results of the evaluation of our deterministic baseline across all folding baselines are shown in Figure 2. We observe, that even for our deterministic design algorithm, the performance strongly depends on the choice of the folding algorithm. Interestingly, the best performing baseline on the folding benchmark, SPOT-RNA, can hardly handle the predictions of our RNA design baseline, resulting in the worst performance. Clearly, our deterministic baseline describes a corner case for structure prediction, since it completely ignores U nucleotides. However, a folding algorithm should be capable of folding any sequence, even if they only consist of G, C, and A nucleotides, so there is still room for improvement.

**Figure 2.**
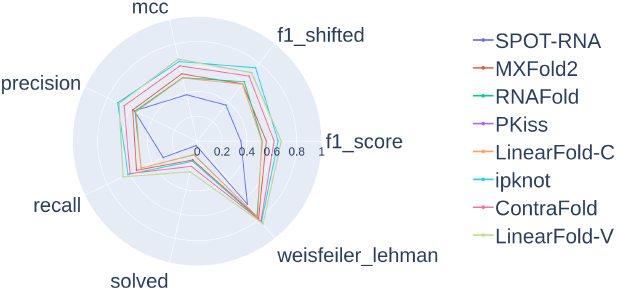
Evaluation of the deterministic baseline for RNA design on inverse RNA folding tasks across all folding base-lines.

### Metric Comparison

We plot the secondary structure of a 5SrRNA of *Drosophila melanogaster* and simulate predictions by shifting all base pairs by one or two positions. The RNA structures are shown in Figure 3. While the structure remains the same, both F1 Score derived measures drop to zero after shifting by one position. Similarly, the MCC drops to a negative value and all three measures stay at these values after another shift by one position. The Weisfeiler-Lehman kernel on the other hand captures the shifting well and the resulting scores successively decrease.

**Figure 3.**
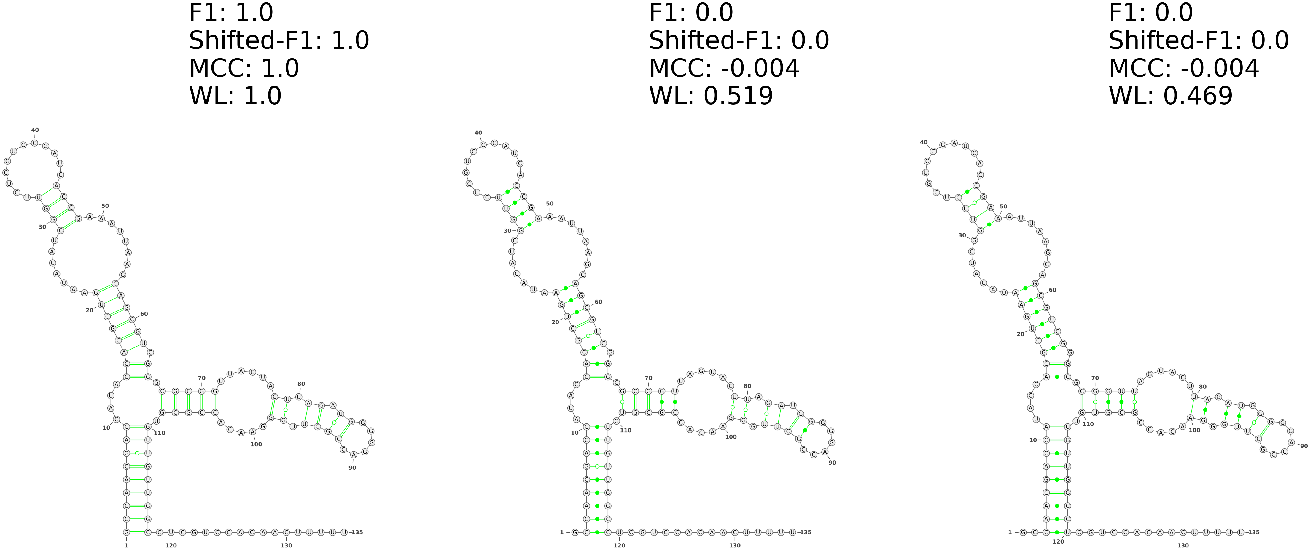
Example of different performance measures using a 5SrRNA of *Drosophila melanogaster*. The original 5SrRNA (left) with all base pairs shifted by one (middle) and two (right) positions. Scores are computed against the original 5SrRNA.

## 5 Discussion

We present RnaBench (RnB), a comprehensive RNA benchmarking library. RnB was developed to standardize the evaluation of algorithms in the field of RNA secondary structure prediction and RNA design as a response to a lack of benchmarks and issues with data curation in the field. In particular, we propose a new data pipeline to split datasets based on biological homology and provide carefully curated training validation and test sets to enable out-of-the-box training and evaluation of algorithms. Additionally, we propose new performance measures to improve the comparability of algorithms. The result is a standardized benchmark for *de novo* RNA secondary structure prediction and the first RNA design benchmark. Moreover, we propose a novel Riboswitch design task geared towards generative algorithms, inspired by recent progress in the field of cheminformatics. Finally, we demonstrate that RnB’s comprehensive visualization module supports the analysis of predictions and helps to capture the strengths and weaknesses of different algorithms, leading to a better understanding and interpretability of the results. However, there are also limitations to our work. Generally, high-quality RNA structure data is rare. While we try to gather as much experimentally supported structure data as possible, still most of the secondary structure data is derived from comparative sequence analysis. However, the generation of high-quality data is out of the scope of this work. Secondly, RNA design is fundamentally linked to RNA folding. A design algorithm can only be as good as the underlying folding engine that folds the designed candidate. We provide a range of folding algorithms to address this issue, however, novel solutions to RNA design that alleviate the dependency are desirable. Nevertheless, we believe that RnaBench is a useful tool to establish deep learning algorithms in the field of RNA computational biology and to support and boost research in the field.

## Appendix

RNA is one of the major regulatory molecules inside living cells and has been connected to diseases like cancer [Prensner et al., 2011] and Parkinson’s [Cao et al., 2018]. While more than 70% of the human genome is transcribed into RNA, only roughly 2% of the genome encodes for proteins [ENCODE Project Consortium and others, 2004, Consortium et al., 2012]. Thus, there is a large number of RNAs that are transcribed from DNA but do not code for proteins. The functions of these so-called *non-coding RNAs* (ncRNAs) remain largely unknown. However, understanding RNA-based regulatory networks is of great interest in fields like biotechnology, synthetic biology, and medicine. Computational RNA structure prediction and design could have a key role to support research as well as experimental screening approaches and boost research in the field. With RnaBench, we present an interface that supports the development of novel algorithms by providing standardized benchmarks, datasets, and utilities that enable out-of-the-box applications.

The supplementary material is structured as follows: In Section A, we report technical details and external software that we use, before describing all our data in detail in Section B. We then detail the performance measures used in RnaBench in Section C, describe the baselines in Section D, and demonstrate the API in Section E.

### A Technical Details and Code Availability

We use the following external software in RnaBench.

#### Baselines

For secondary structure prediction, we use RNAFold from the ViennaRNA package [Lorenz et al., 2011] version 2.5.0, LinearFold [Huang et al., 2019] version 1.0, ContraFold [Do et al., 2006] version 2.02 as provided via anaconda, SPOT-RNA [Singh et al., 2019] downloaded from https://github.com/jaswindersingh2/SPOT-RNA, MXFold2 [Sato et al., 2021] version 0.1.2, IPknot [Sato et al., 2011] version 1.1.0, and pKiss [Janssen and Giegerich, 2015] version 2.2.14 provided by anaconda. For RNA design, we use RNAInverse version 2.5.0 as provided by the ViennaRNA package.

#### Data

For our data pipelines, we use BLAST-N [Altschul et al., 1997] version 2.12.0, Infernal [Nawrocki and Eddy, 2013] version 1.1.4, bpRNA [Danaee et al., 2018], downloaded via https://github.com/hendrixlab/bpRNA, CD-HIT [Fu et al., 2012] version 4.8.1, LocARNA-P [Will et al., 2012] version 2.0.0RC10, and get 3D data from RNAsolo [Adamczyk et al., 2022] with BGSU version 3.286 as described at http://rna.bgsu.edu/rna3dhub/nrlist.

#### Utilitites

For different low-level utilities, we mainly depend on Biopython [Cock et al., 2009]. We use forgi [Thiel et al., 2019] version 2.1.2 for extracting secondary structure motifs. For plotting molecular structures, we use Varna [Darty et al., 2009] version 3.9.

Our benchmark package and source code are attached to the supplementary material and will be open-sourced with the acceptance of this paper.

### B Data

In this section, we detail all datasets that we use for RnaBench. However, we start with an explanation of the different RNA secondary structure representations as shown in Figure 4 in Section B.1. Then, we describe the datasets and databases that we used to build our initial training data pool of 111,295 samples in Section B.2. In Section B.3, we describe the test sets used for the RNA secondary structure prediction benchmarks, and in Section B.4, we detail the test sets used for our RNA design benchmark.

**Figure 4.**
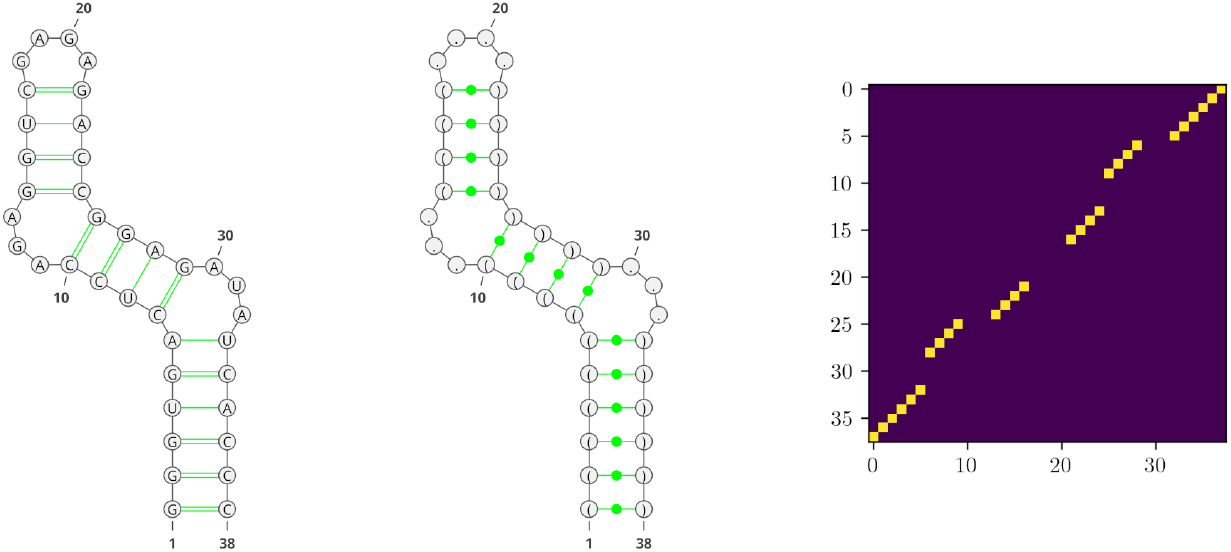
Three representations of the same RNA secondary structure. (Left) Common graph representation of the RNA. (Middle) We display the dot-bracket notation in the graph structure. Paired nucleotides are indicated as a pair of matching brackets, unpaired nucleotides are indicated by a dot. (Right) Matrix representation of the RNA. This so-called contact map is a binary *L × L* square matrix, where *L* is the sequence length of the RNA. Pairing nucleotides are shown in yellow.

#### B.1 RNA Secondary Structure Representations

RNA sequences are described with the four nucleobases **A**denine, **C**ytosine, **G**uanine, and **U**racil2 (A, C, G, U; commonly referred to as nucleotides). While RNA 3D structures are typically represented as a list of (*x, y, z*) atom coordinates, RNA secondary structures can be viewed as a graph, where nodes correspond to nucleotides of the RNA sequence and edges describe connections between nucleotides that form base-pairs via hydrogen bonds. From the computational point of view, these graphs can be represented in different ways e.g. using string notations [Hofacker et al., 1994] (Figure 4, middle), binary adjacency matrices (contact maps) of size *L × L*, where *L* is the length of the RNA sequence (Figure 4, right), or a list of pairs that indicates the indices of interacting nucleotides of the RNA sequence. The most common types of base pairs are formed between A and U, G and C, or G and U, known as canonical pairs, but all other combinations are possible and have been reported. Depending on the requirements of the application, the structure representations have different advantages. String notations are simple and allow to indicate different levels of nesting, i.e. pseudoknots [Staple and Butcher, 2005], in the structure via different types of brackets. However, While most nucleotide interactions are formed between two pairing partners, substructures might be formed by interactions of more than two nucleotides (base triples, base multiplets). The common string notations cannot describe these types of interactions, while contact maps can. Due to the binary nature of the adjacency matrix representation, however, these lack pseudoknot information. Therefore, we represent RNA secondary structures as lists of pairs in RnaBench, but additionally, add the pseudoknot information to the pairs. This allows us to get the best of both worlds. We display different types of secondary structure representations in Figure 4.

#### B.2 Datasets of the Training Data Pool

In this section, we detail the datasets and databases that we use for the creation of the initial training data pool. For all datasets, we use only sequences that do not contain any gap symbols.

##### bpRNA-1m Meta-Database

The bpRNA-1m meta-database [Danaee et al., 2018] is a publicly available collection of annotated RNA sequences and structures, comprising 102,318 samples. The data is gathered from the following public databases: the Comparative RNA Web (CRW) [Cannone et al., 2002], tmRNA database [Zwieb et al., 2003], tRNAdb [Andersen et al., 2006], Signal Recognition Particle (SRP) database [Larsen et al., 1998], RNase P database [Brown, 1998], tRNAdb 2009 database [Jühling et al., 2009], all data from the RCSB Protein Data Bank (PDB) [wwp, 2019] that consists of one RNA molecule as of 12 June 2017, and all families from the RNA Family database (Rfam) [Griffiths-Jones et al., 2003], version 12.2. The annotations in bpRNA-1m include pseudoknots [Staple and Butcher, 2005] and non-canonical base pairs.

##### ArchiveII

We use the ArchiveII dataset as provided by Chen et al. [2020], that was originally proposed by Sloma and Mathews [2016]. The ArchiveII dataset consists of 3975 samples from the following RNA families: small subunit ribosomal RNA [Gutell, 1994], large subunit ribosomal RNA [Gutell et al., 1993, Schnare et al., 1996], 5S ribosomal RNA [Szymanski et al., 1998, Daub et al., 2008], Group I self-splicing introns [Waring and Davies, 1984, Damberger and Gutell, 1994], RNase P RNA [Brown, 1998], signal recognition particle RNA [Larsen et al., 1998], tRNA [Sprinzl et al., 1998], and tmRNA [Zwieb and Wower, 2000].

##### RNAStralign

RNAStralign is a database of known homologous sequences and structures. The families included in this database are: 5S ribosomal RNA [Szymanski et al., 2002], Group I intron [Zhou et al., 2008], tmRNA [Zwieb et al., 2003], tRNA [Jühling et al., 2009], 16S ribosomal RNA [Cannone et al., 2002], Signal Recognition Particle (SRP) RNA [Gorodkin et al., 2001], RNase P RNA [Brown, 1998] and telomerase RNA [Nawrocki et al., 2015]. Our version of RNAStralign contains 37,136 samples.

##### RNA-Strand

RNA-Strand [Andronescu et al., 2008] is a database of 4666 RNAs with known secondary structures of any type and organism. RNA-Strand is available at http://www.rnasoft.ca/strand/.

##### Datasets provided by

**Sato et al**. **[2021]** We use the following datasets provided by Sato et al. [2021]: The TrainSetA, TestSetA, TrainSetB, TestSetB [Rivas et al., 2012], and the bpRNA-new set [Sato et al., 2021].

##### Rfam-Learn Dataset

We also gathered the Rfam-Learn dataset provided by [Runge et al., 2019]. However, we only report this dataset for completeness, since we dropped the entire dataset during data preparation because the RNA sequences of this set are already masked due to the task of inverse RNA folding that was tackled by Runge et al. [2019].

##### Dataset provided by

**Taneda [2010]** We also include the dataset provided by Taneda [2010]. This dataset contains 29 samples.

##### Eterna100

We include the Eterna100 dataset version 1 [Anderson-Lee et al., 2016] and 2 [Koodli et al., 2021]. We use the first solution for each RNA structure if available. The datasets were downloaded from https://github.com/eternagame/eterna100-benchmarking.

##### Datasets provided by

**Kleinkauf et al**. **[2015]** We use all training sets as provided by Kleinkauf et al. [2015]. The datasets were downloaded from https://github.com/RobertKleinkauf/antarna/tree/antaRNAdp/ Constraints. In particular, we use the Pseudobase++ [Taufer et al., 2009] train set and the Rfam [Griffiths-Jones et al., 2003] train set for our initial pool.

##### Protein Data Bank

We download all RNA containing samples from PDB [wwp, 2019] in May, 2022. We process the 3D structures using DSSR [Lu et al., 2015] and obtain structure annotations using bpRNA [Danaee et al., 2018]. The dataset contains a total of 13,334 samples.

#### B.3 Secondary Structure Prediction Benchmark Test and Validation Datasets

For our secondary structure prediction benchmark, we use the validation and test datasets provided by Singh et al. [2019] and Singh et al. [2021]. For validation, we use VL0 and VL1, provided by Singh et al. [2019]. Our versions of these datasets contain 1300 and 29 samples, respectively. VL0 is derived from the bpRNA meta-database and VL1 is derived from PDB samples at a resolution threshold of 3.5Å. We now detail the different test sets TS0, TS1, TS2, TS3, and TS-hard.

##### TS0

The TS0 dataset is derived from the bpRNA meta-database and consists of 1305 samples. We use the TS0 dataset provided by Singh et al. [2019].

##### TS1

We use the TS1 dataset provided by Singh et al. [2019]. The dataset consists of 67 samples derived from high-resolution (<3.5Å) 3D RNA structures from PDB, downloaded in March 2019.

##### TS2

The TS2 dataset is derived from NMR structures of the PDB and consists of 39 samples. Again we use the original version of this set as provided by Singh et al. [2019].

##### TS3

The TS3 dataset we use here is provided by Singh et al. [2021] and contains 19 samples. The dataset was derived from PDB 3D structure data downloaded in April 2020.

##### TS-Hard

The testset TS-hard is derived from the high-resolution datasets TS1 and TS3 and contains 28 samples. Originally, this dataset was created by Singh et al. [2021].

###### B.3.1 Secondary Structure Prediction Benchmark Dataset Overview

We show the length distributions of all datasets of the RNA secondary structure benchmarks. Figure 5 shows the datasets for the inter-family prediction benchmark, Figure 6 for the intra-family prediction benchmark, and Figure 7 shows the length distribution of the datasets of the biophysical model benchmark. We observe that the length distribution of our inter-family benchmark differs between the training, validation and test splits, while these sets are more similar for the other two benchmarks. We mainly attribute this to the more strict pipeline for removing data homologies for the inter-family benchmark compared to the other two benchmarks and again note the importance of carefully curated datasets that account for homologies.

**Figure 5.**
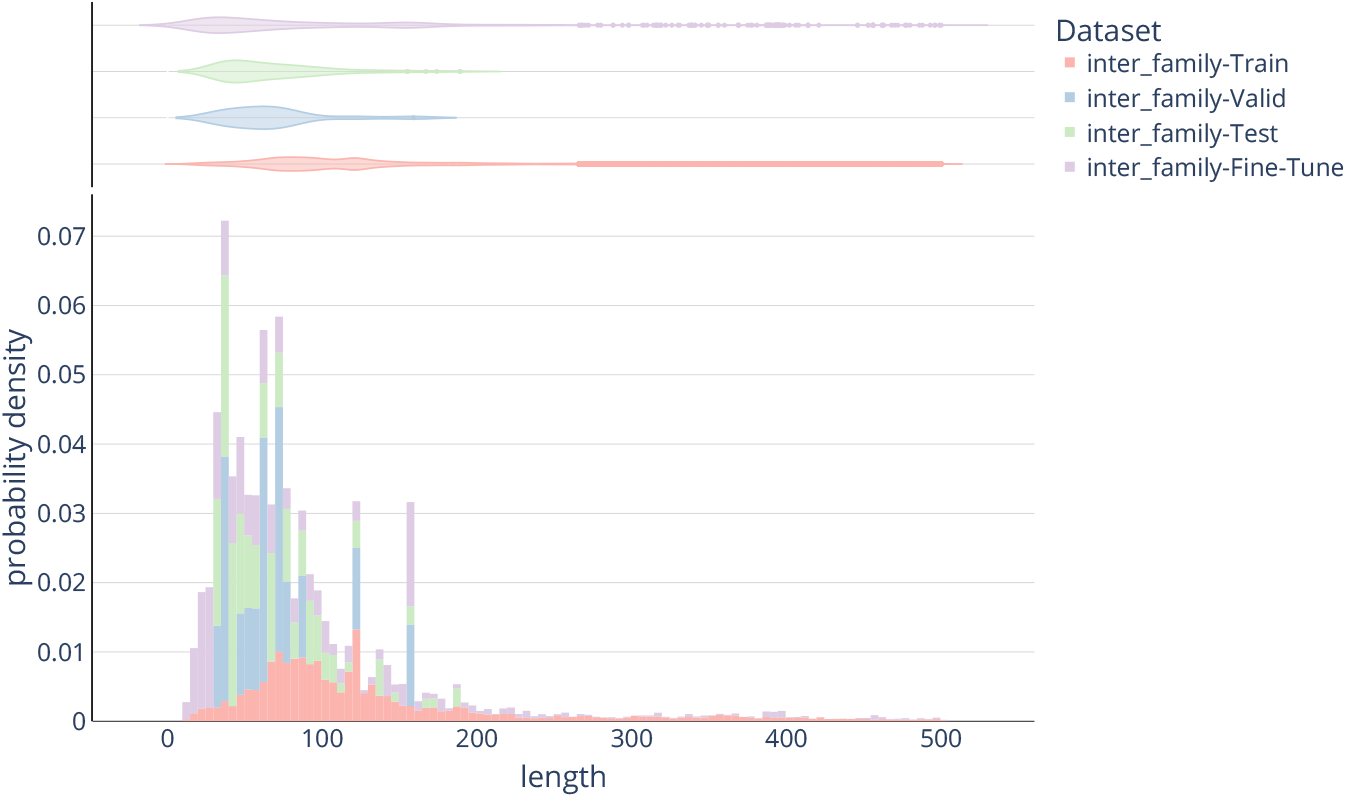
Length distribution of all datasets of the Inter-Family Benchmark.

**Figure 6.**
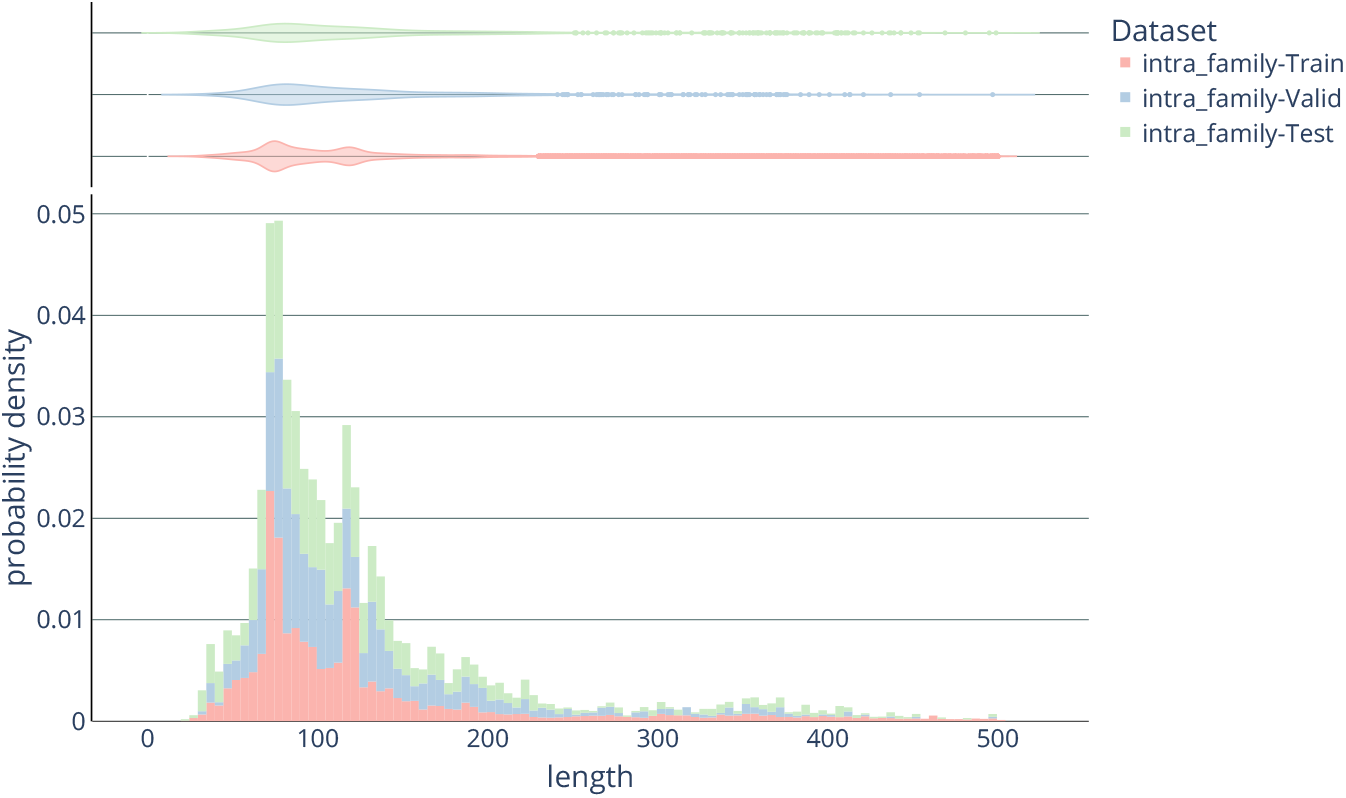
Length distribution of all datasets of the Intra-Family Benchmark.

**Figure 7.**
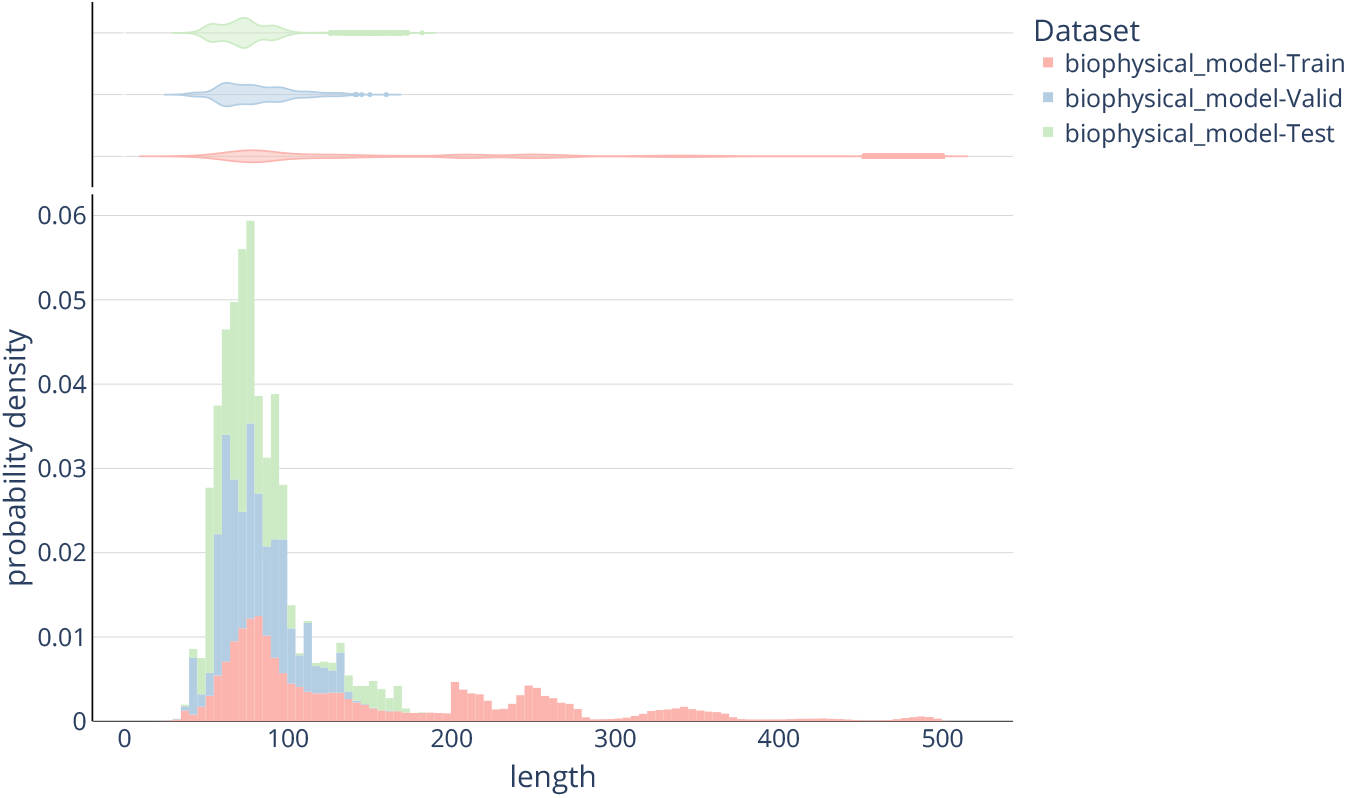
Length distribution of all datasets of the Biophysical Model Benchmark.

#### B.4 RNA Design Benchmark Test Datasets

For the design benchmark, we use different test sets from the literature. The validation data is sampled from the training set after ensuring that there is no overlap between the test and training sets. We now detail the different test sets.

##### Rfam-Learn

The Rfam-Learn dataset was created by Runge et al. [2019] and contains 100 samples. The dataset was derived from sequences of the Rfam database version 13.0. and structures are predicted structures using RNAFold [Lorenz et al., 2011].

##### Rfam Testset provided by

**Taneda [2010]** The testset provided by Taneda [2010] contains 29 samples. The dataset was derived from the Rfam database version 9.0.

##### RNA-Strand set provided by

**Kleinkauf et al**. **[2015]** The RNA-Strand dataset provided by Kleinkauf et al. [2015] contains a total of 50 samples. We download the dataset from https://github.com/RobertKleinkauf/antarna/tree/antaRNAdp/Constraints.

##### ArchiveII

We use all pseudoknot containing RNAs from the ArchiveII dataset.

##### Pseudobase++ Testset

The pseudobase++ dataset provided by Kleinkauf et al. [2015] contains 252 samples and can be downloaded via https://github.com/RobertKleinkauf/antarna/tree/antaRNAdp/Constraints.

##### Rfam Testset provided by

**Kleinkauf et al**. **[2015]** The Rfam testset provided by Kleinkauf et al. [2015] consists of 63 samples that can be donloaded from https://github.com/RobertKleinkauf/antarna/tree/antaRNAdp/Constraints.

##### Eterna100

Koodli et al. [2021] recently showed that the original version of the Eterna100 test set contains 19 samples that are unsolvable with the set of thermodynamic **parameters** implemented in the ViennaRNA package **[Lorenz et al**., **2011]** version 1. We, therefore, use two versions of the Eterna100 dataset, version 1 as provided by **Anderson-Lee et al**. **[2016]** and version 2 as provided by **Koodli et al**. **[2021]**. Both datasets can be downloaded from https://github.com/eternagame/eterna100-benchmarking. The datasets contain 100 samples each.

###### B.4.1 RNA Design Benchmark Dataset Overview

We show the length distribution of the inverse RNA folding benchmark datasets in Figure 8 and for the constrained-design benchmark in Figure 9. For the inverse RNA folding benchmark, we observe the largest difference between training and test data, with a cluster of samples with a length between 300 and 400 nucleotides in the test data. These samples are mainly coming from the ArchiveII dataset, which contains longer samples than the other test sets.

**Figure 8.**
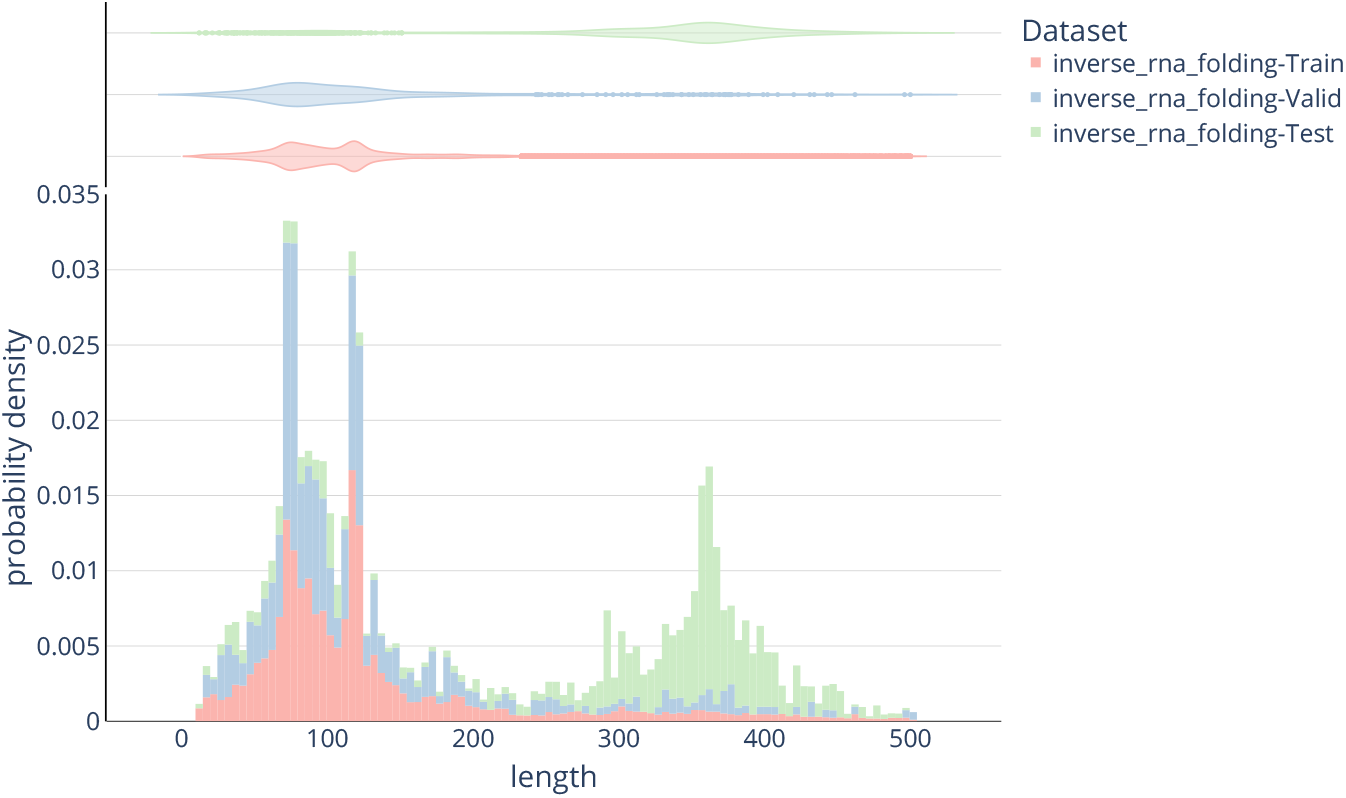
Length distribution of all datasets of the Inverse RNA Folding Benchmark.

**Figure 9.**
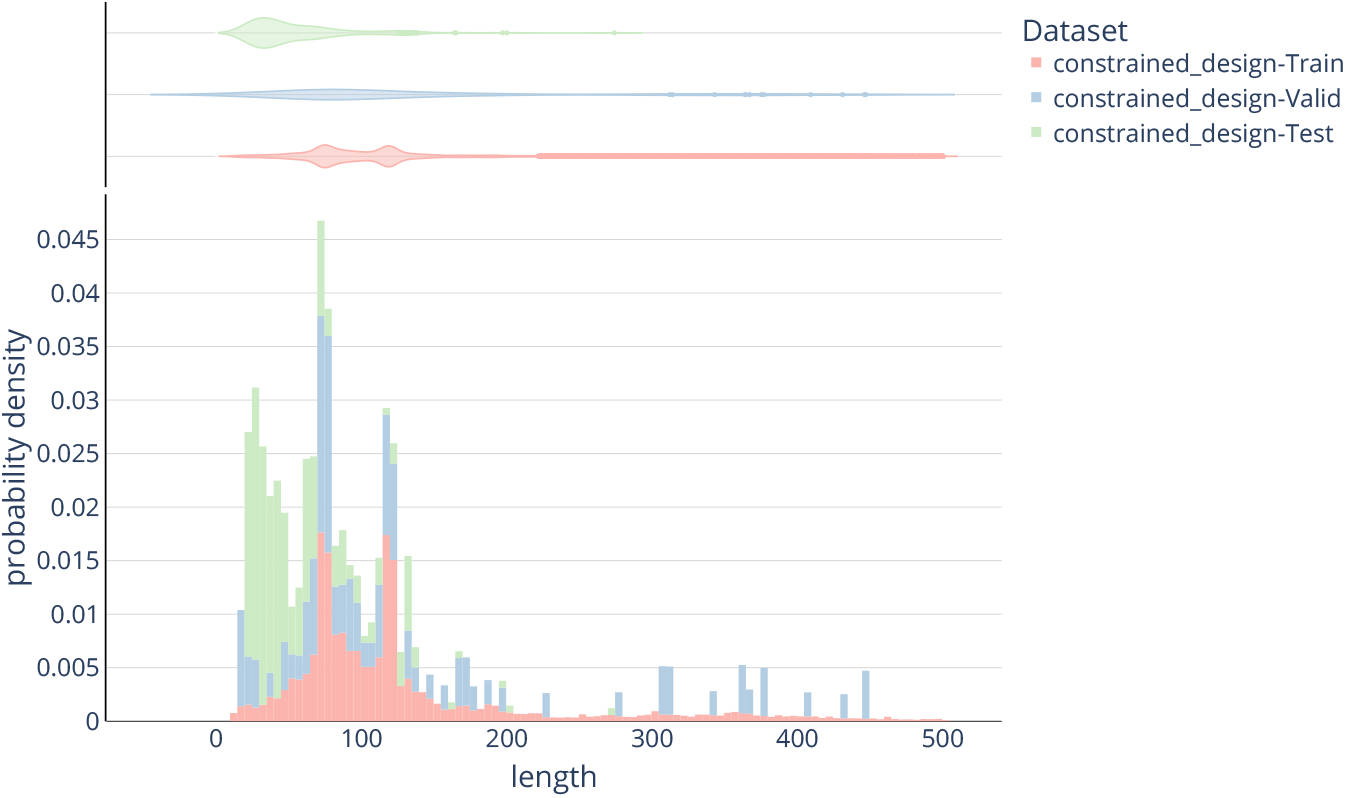
Length distribution of all datasets of the Constrained Design Benchmark.

#### B.5 3D Data

RnaBench also includes pipelines for downloading, parsing, and loading 3D data. For our 3D data, we use RNA-solo [Adamczyk et al., 2022], a repository of cleaned RNA 3D structures from PDB. The cleaning procedure includes removing protein structures from the files as well as dividing RNAs into equivalence classes following the BGSU guidelines. RNAsolo further provides RNA 3D structures at different resolution cutoffs, which can be chosen in the download script.

### C Evaluation

For evaluation, we use commonly used and novel performance measures, described in the following. The commonly used performance measures for RNA secondary structure prediction are based on a confusion matrix, which describes the number of true positives (TP), true negatives (TN), false positives (FP), and false negatives (FN) of a given prediction.

#### Binary Classification Measures

We use three standard measures:

#### Precision

describes the fraction of correctly predicted base pairs and is calculated as

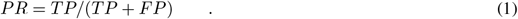

#### Recall

or sensitivity describes the true positive predictions, calculated as

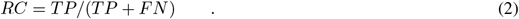

The **F1 Score** is the harmonic mean of sensitivity and precision

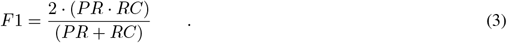

#### Shifted F1 Score

Recent work in the field of secondary structure prediction proposed to use a shifted version of the F1 Score to account for the dynamic nature of RNA [Mathews, 2019], in particular bulge migration events [Woodson and Crothers, 1987]. The shifted F1 Score is calculated as the F1 score, but for a given pair (*i, j*) all pairs (*i, j* + 1), (*i* + 1, *j*), (*i, j −* 1), and (*i −* 1, *j*) are also considered correct.

#### Matthews Correlation Coefficient (MCC)

While the F1 score emphasizes positives, the MCC is a more balanced measure. The MCC can be calculated as follows.

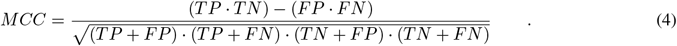

#### Weisfeiler-Lehman (WL) Graph Kernel

The WL kernel is a powerful metric for comparing graphs, as it captures the structural information while being computationally efficient. It has been widely adopted in graph classification, pattern recognition, and graph mining tasks, showcasing its effectiveness across various domains.

#### Solved Structures

While it is the ultimate goal of secondary structure prediction, publications in the field typically do not report the number of completely correct predicted structures.

We further allow analyzing predictions concerning different measures that assess the learning behavior from a distribution learning point of view. In particular, we provide the following measures.

#### Intersection Over Union Complement IoUC

between two sets *G* and *O* is given by 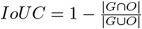 as a measure of novelty.

#### Novelty

The difference between IOUC metric and Novelty metric lies in the denominator of the division: IoUC calculates the ratio of the intersection |*G ∩ O*| to the union of the generated set *G* and training set |*O*|, while Novelty calculates the ratio of the intersection to the size of the generated set |*G*| itself.

#### Diversity

Internal Diversity and Diameter are two key measures used to assess the diversity of generated RNA structures. These measures rely on a self-distance matrix *d*, referred to as the pairwise distance matrix, computed between each pair of elements in the generated dataset according to an arbitrary distance measure. Diversity captures the average difference within the pairwise distance matrix while excluding the distance of an element to itself. On the other hand, Diameter [Edelsbrunner and Harer, 2010] represents the maximum difference observed in the pairwise distance matrix.

#### Inference time

RnaBench further provides per-sample timing, while the mean runtime is provided as an additional performance measure.

##### C.1 Riboswitch Design

For the riboswitch design, we reimplement the evaluation procedure of Wachsmuth et al. [2012]. In particular, all designed candidate sequences are validated based on features of the sequence and the structure of the constructs. These features include: A shape of exactly two hairpins, the existence of the exact aptamer hairpin structure, the formation of the terminator hairpin between the 3^*′*^-end of the aptamer, the spacer and the region complementary to the aptamer sequence, no pairing within the last seven nucleotides of the 8-U-stretch, and no pairing between the spacer and the aptamer in a sequence of folding steps, simulating co-transcriptional folding with a fixed elongation speed of five. Further, we assess the predicted sequences for the existence of the aptamer sequence, a minimum size of four unpaired nucleotides in the spacer region, and the sequence of the 8-U stretch.

Additionally, as part of the generative assessment, we evaluate the novelty and diversity of the generated sequences and their corresponding structure pairs (see Table 2). We also compute the KL divergence between the original data and the generated data for the Riboswitch design based on a set of selected features such as structural motifs, such as hairpins, stems, and internal loops. (see Table 3) This comprehensive evaluation provides insights into the structural integrity, novelty, and diversity of the designed RNA sequences.

**Table 3:**
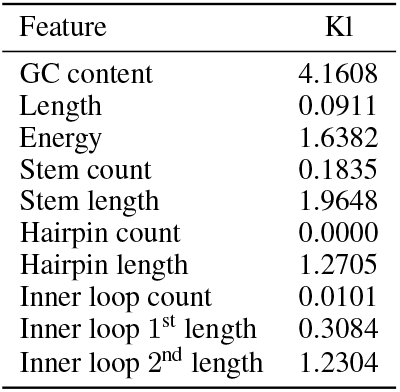
KL Divergence between the train and generated Data for Riboswitch Design.

### D Baselines

We use selected baselines for all benchmarks in RnaBench.

#### D.1 RNA Secondary Structure Prediction

For the secondary structure prediction benchmark, we try to provide a broad range of different approaches with RnaBench.

##### RNAFold

RNAFold [Hofacker et al., 1994] is one of the most commonly used secondary structure prediction algorithm, provided with the ViennaRNA package [Lorenz et al., 2011]. We use version 2.5.0 of RNAFold in RnaBench, which uses the thermodynamic parameters of the 2004 Turner model [Mathews et al., 2004]. RNAFold applies an energy minimization approach to find the most stable secondary structure, called the minimum free energy structure.

##### ContraFold

ContraFold [Do et al., 2006] is an algorithm that uses a Nussinov-like recursion [Nussinov et al., 1978] to predict the secondary structure with the maximum expected accuracy (MEA), using McCaskill’s algorithm [McCaskill, 1990].

##### LinearFold

LinearFold [Huang et al., 2019] uses a linear-time approximation of the partition function to predict secondary structures. We use both versions of LinearFold, LinearFold-V based on the ViennaRNA folding engine, and LinearFold-C based on the ContraFold engine.

##### pKiss

pKiss [Janssen and Giegerich, 2015] is the successor of pknotsRG [Reeder and Giegerich, 2004] and can predict two limited classes of pseudoknots using heuristics.

##### Ipknot

IPknot [Sato et al., 2011] predicts secondary structures with pseudoknots by maximizing expected accuracy of the predicted structure. IPknot decomposes a pseudoknotted structure into a set of pseudoknot-free substructures and approximates base pair probability distributions that consider pseudoknots.

##### SPOT-RNA

SPOT-RNA Singh et al. [2019] was the first algorithm using deep neural networks for end-to-end prediction of RNA secondary structures, using an ensemble of models with residual networks (ResNets) He et al. [2016], bidirectional LSTM-[Hochreiter and Schmidhuber, 1997] (BiLSTMs) [Schuster and Paliwal, 1997], and dilated convolution [Yu and Koltun, 2015] architectures. *SPOT-RNA* was trained on a large set of intra-family RNA data for predictions on TS0, and further fine-tuned on a small set of experimentally-derived RNA structures.

##### MXFold2

MXFold2 Sato et al. [2021] combines deep learning with a Dynamic Programming (DP) approach using a CNN/BiLSTM architecture to learn the scoring function for the DP algorithm. The network is trained to predict scores close to a set of thermodynamic parameters to increase robustness. MXFold2 is restricted to predicting a reduced set of base pairs due to limitations in the DP algorithm.

##### D.2 RNA Design

RNA design algorithms are typically depending on a specific secondary structure prediction algorithm. To remain mostly independent of the folding algorithm, we choose to implement a simple solution to the RNA design problem. However, we also include RNAInverse [Hofacker et al., 1994] as a commonly used inverse RNA folding algorithm.

##### RNAInverse

RNAInverse [Hofacker et al., 1994] is a commonly used inverse RNA folding algorithm from the ViennaRNA [Lorenz et al., 2011] package.

##### Deterministic Baseline

We implement a design algorithm that is independent of the folding engine. Our deterministic baseline predicts an A-nucleotide for each position that is unpaired and a G-C base pair for all positions ought to be paired.

#### D.3 Riboswitch Design

As described in the main part of this work, we reimplement the original procedure of Wachsmuth et al. [2012] as a baseline for our Riboswitch design benchmark.

##### E. API

We provide a simple interface for RnaBench as shown in the code example 1. To interact with RnaBench, define a function that wraps your model predictions. This function then is evaluated in the benchmark. Depending on the benchmark task, the returned dictionary contains a different set of performance measures to assess the performance of the algorithm.

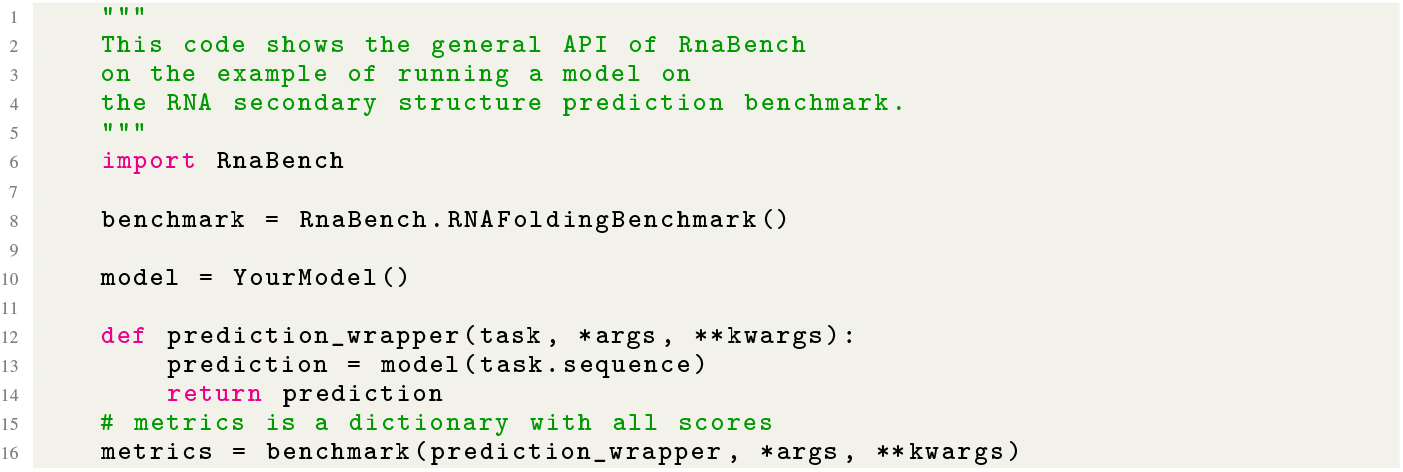

Code 1: Example API.

We also provide interfaces to get the data provided with a PyTorch [Paszke et al., 2019] data loader. The code example 2 shows an example.

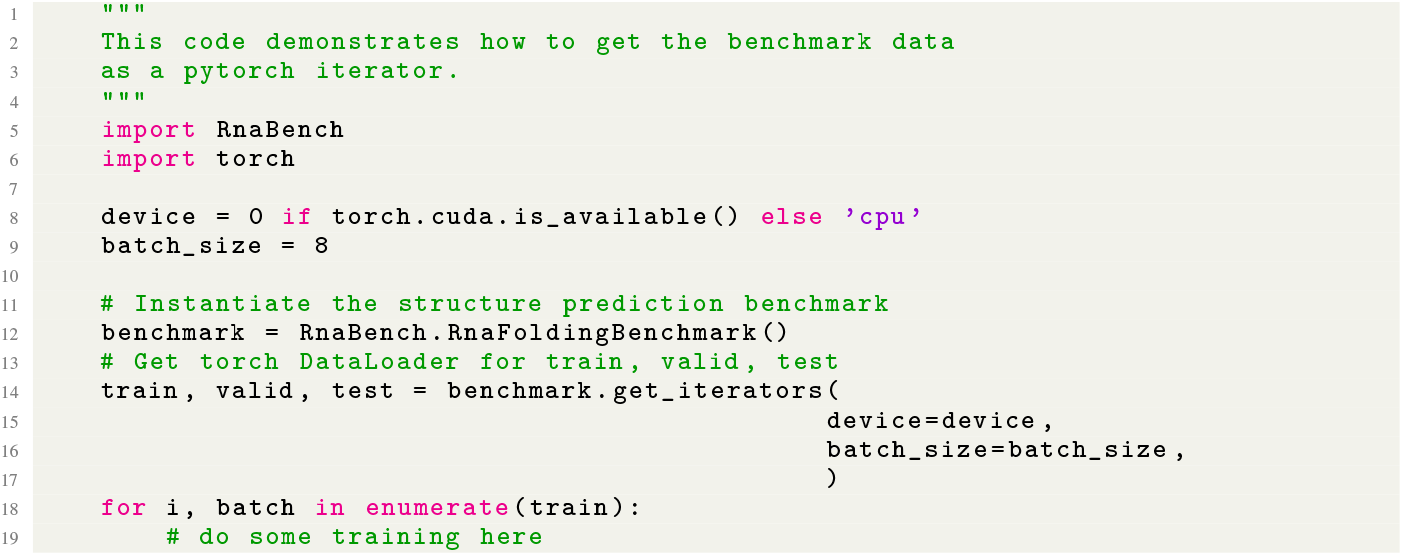

Code 2: Usage example with a pyTorch DataLoader.

We provide a range of further examples with RnaBench as well as scripts to reproduce our experiments.

## Footnotes

1 Our code is publicly available via addsomeurlhere.

2 Extensions are known e.g. to represent modified nucleotides as described at the Ligand Expo website hosted by the RCSB PDB at http://ligand-expo.rcsb.org/index.html, or using the IUPAC nomenclature described by the International Nucleotide Sequence Database Collaboration (INSDC) at https://www.insdc.org/documents/feature_table.html#7.4.1.

## Notes

https://pypi.org/project/RnaBench/

### Competing Interest Statement

The authors have declared no competing interest.

